# Ipomoeassin-F inhibits the *in vitro* biogenesis of the SARS-CoV-2 spike protein and its host cell membrane receptor

**DOI:** 10.1101/2020.11.24.390039

**Authors:** Sarah O’Keefe, Peristera Roboti, Kwabena B. Duah, Guanghui Zong, Hayden Schneider, Wei Q. Shi, Stephen High

## Abstract

In order to produce proteins essential for their propagation, many pathogenic human viruses, including SARS-CoV-2 the causative agent of COVID-19 respiratory disease, commandeer host biosynthetic machineries and mechanisms. Three major structural proteins, the spike, envelope and membrane proteins, are amongst several SARS-CoV-2 components synthesised at the endoplasmic reticulum (ER) of infected human cells prior to the assembly of new viral particles. Hence, the inhibition of membrane protein synthesis at the ER is an attractive strategy for reducing the pathogenicity of SARS-CoV-2 and other obligate viral pathogens. Using an *in vitro* system, we demonstrate that the small molecule inhibitor ipomoeassin F (Ipom-F) potently blocks the Sec61-mediated ER membrane translocation/insertion of three therapeutic protein targets for SARS-CoV-2 infection; the viral spike and ORF8 proteins together with angiotensin-converting enzyme 2, the host cell plasma membrane receptor. Our findings highlight the potential for using ER protein translocation inhibitors such as Ipom-F as host-targeting, broad-spectrum, antiviral agents.

## Introduction

Many viruses, including SARS-CoV-2 (Zhou et al., 2020; Zhu et al., 2020) (Fig. 1A), hijack the host cell secretory pathway to correctly synthesise, fold and assemble important viral proteins (Bojkova et al., 2020; Gordon et al., 2020; Sicari et al., 2020). Hence, small molecule inhibitors of Sec61-mediated co-translational protein entry into the endoplasmic reticulum (ER) (Luesch and Paavilainen, 2020) have potential as broad-spectrum antivirals (Heaton et al., 2016; Shah et al., 2018). Such inhibitors offer a dual approach; first, by directly inhibiting production of key viral proteins and, second, by reducing levels of host proteins co-opted during viral infection. Hence, human angiotensin-converting enzyme 2 (ACE2) is an important host cell receptor for SARS-CoV-2 viral entry (Cantuti-Castelvetri et al., 2020; Daly et al., 2020; Walls et al, 2020) synthesised at the ER prior to its trafficking to the plasma membrane (Warner et al., 2005).

**Fig. 1.**
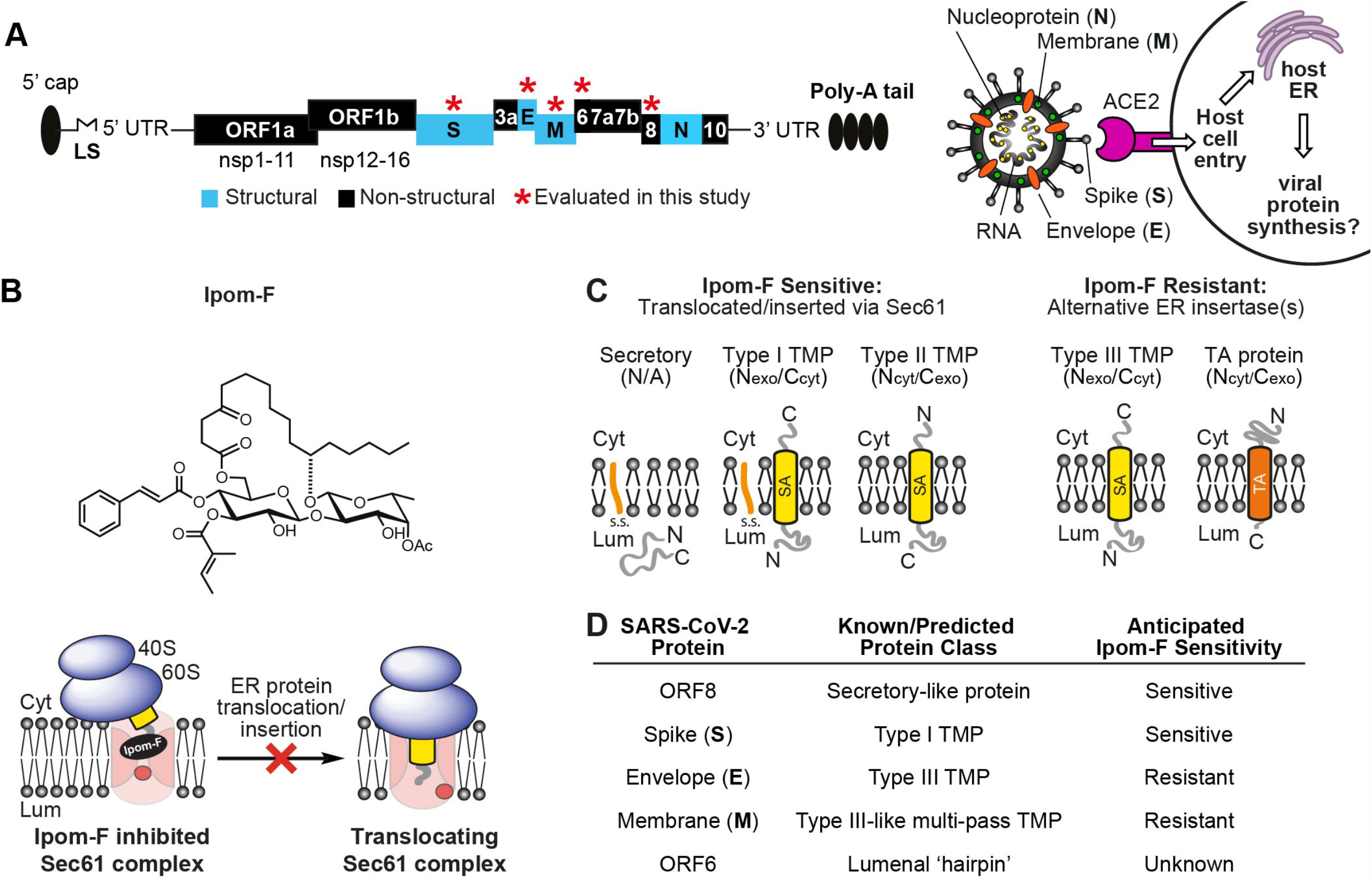
Ipom-F as a potential inhibitor of SARS-CoV-2 viral protein synthesis. **(A)** Schematic of (+) ssRNA genome architecture of SARS-CoV-2 (29903 nt) containing 5’ capped mRNA with a leader sequence (LS), 3’ end poly-A tail, 5’ and 3’ UTRs and open reading frames (ORFs): ORF1a, ORF1b, spike (S), ORF3a, envelope (E), membrane (M), ORF6, ORF7, ORF8, nucleoprotein (N) and ORF10 (Firth, 2020; Naqvi et al., 2020). An important mode of SARS-CoV-2 host entry proceeds via interaction of the viral S protein with human angiotensin-converting enzyme 2 (ACE2) (Walls et al., 2020). (**B**) Structure of Ipomoeassin-F (Ipom-F), a small molecule inhibitor of Sec61-mediated protein translocation. (**C**) Ipom-F efficiently blocks membrane translocation of secretory proteins and insertion of single-pass type I and type II TMPs, but not insertion of type III TMPs or tail-anchored (TA) proteins. SA denotes a signal anchor. (**D**) Based on known/predicted membrane topology of SARS-CoV-2 proteins, and sensitivity of comparable host cell proteins (Zong et al., 2019; O’Keefe et al., 2020 submitted), likely sensitivity to Ipom-F was anticipated.

Our recent studies show that ipomoeassin-F (Ipom-F) (Fig. 1B) is a potent and selective inhibitor of Sec61-mediated protein translocation at the ER membrane (Zong et al., 2019; O’Keefe et al., 2020 submitted). Given that SARS-CoV-2 membrane proteins likely co-opt host mechanisms of ER entry (cf. Gordon et al., 2020; Sicari et al., 2020), we concluded that their sensitivity to Ipom-F would likely be comparable to that of endogenous Sec61 clients (Fig. 1C; see also Zong et al., 2019; O’Keefe et al., 2020 submitted). We, therefore, evaluated the effects of Ipom-F on SARS-CoV-2 proteins containing hydrophobic ER targeting signals (Fig. 1D). The *in vitro* membrane insertion of the viral spike (S) protein and membrane translocation of the ORF8 protein are both strongly inhibited by Ipom-F, whilst several other viral membrane proteins are unaffected (Fig. 2). Likewise, the ER integration of ACE2, an important host receptor for SARS-CoV-2 (Walls et al., 2020), is highly sensitive to Ipom-F (Fig. 2).

**Fig. 2.**
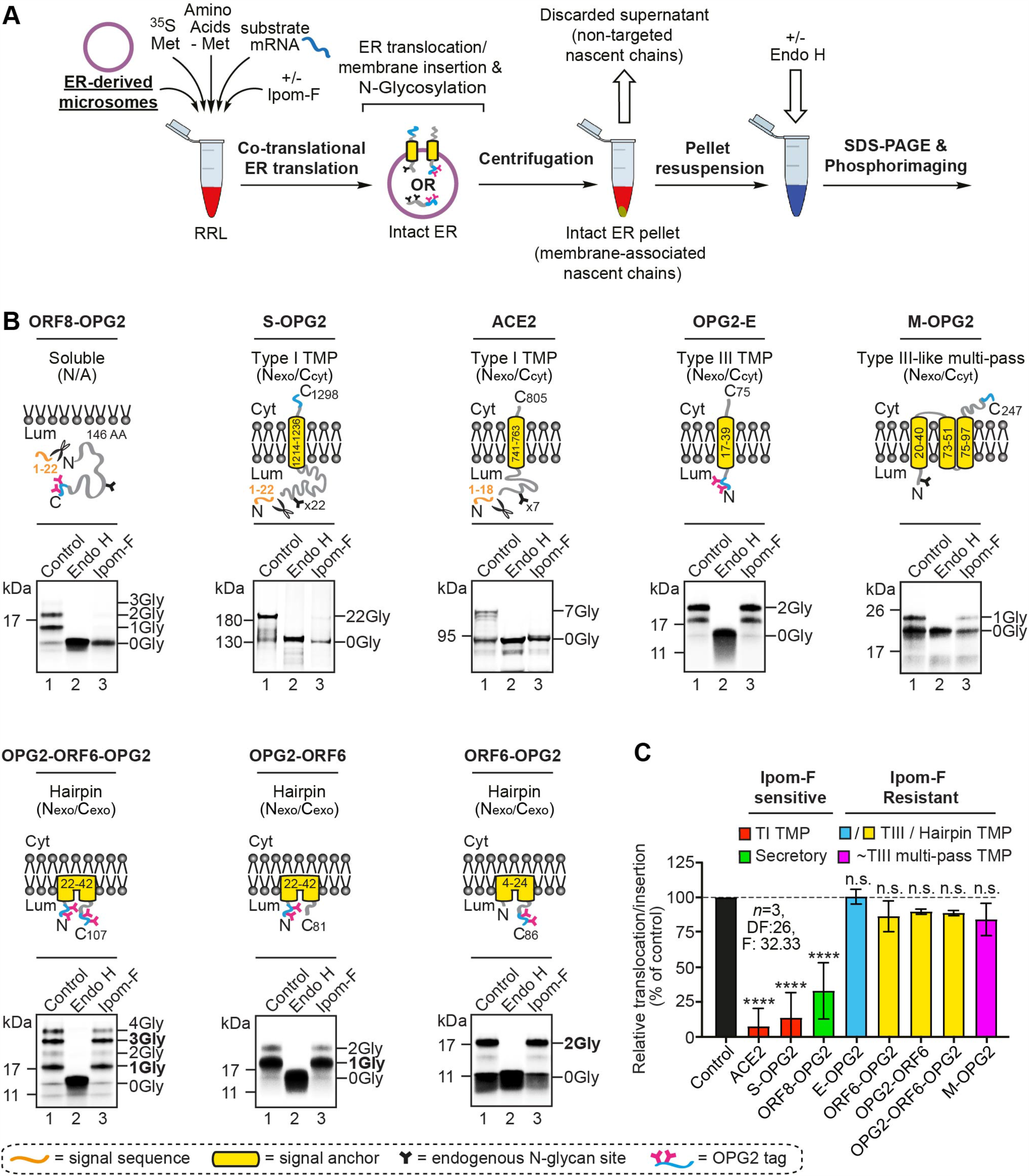
Ipom-F selectively inhibits the ER membrane translocation of SARS-CoV-2 proteins. **(A)** Schematic of *in vitro* ER import assay using pancreatic microsomes. Following translation, fully translocated/membrane inserted radiolabelled precursor proteins are recovered and analysed by SDS-PAGE and phosphorimaging. N-glycosylated species were confirmed by treatment with endoglycosidase H (Endo H). **(B)** Protein precursors of the human angiotensin-converting enzyme 2 (ACE2) and OPG2-tagged versions of the SARS-CoV-2 ORF8 (ORF8-OPG2), spike (S-OPG2), envelope (OPG2-E), membrane (M-OPG2) and ORF6 (a doubly-OPG2 tagged version, OPG2-ORF6-OPG2, and two singly-OPG2 tagged forms, OPG2-ORF6 and ORF6-OPG2, with predominant N-glycosylated species in bold) were synthesised in rabbit reticulocyte lysate supplemented with ER microsomes without or with Ipom-F (lanes 1 and 3). Phosphorimages of membrane-associated products resolved by SDS-PAGE with representative substrate outlines are shown. N-glycosylation was used to measure the efficiency of membrane translocation/insertion and N-glycosylated (X-Gly) versus non-N-glycosylated (0Gly) species identified using Endo H (see lane 2). **(C)** The relative efficiency of membrane translocation/insertion in the presence of Ipom-F was calculated using the ratio of N-glycosylated protein to non-glycosylated protein, relative to the DMSO treated control (set to 100% efficiency). Quantitations are given as mean±s.e.m for independent translation reactions performed in triplicate (*n*=3) and statistical significance (one-way ANOVA, DF and F values shown in the figure) was determined using Dunnett’s multiple comparisons test. Statistical significance: n.s., non-significant >0.1; ****, P < 0.0001.

We show that the principle molecular basis for the Ipom-F sensitivity of SARS-CoV-2 proteins is their dependence on Sec61, as dictated by their individual structural features and membrane topologies (Fig. 3). Taken together, our *in vitro* study of SARS-CoV-2 protein synthesis at the ER highlights Ipom-F as a promising candidate for the development of a broad-spectrum, host-targeting, antiviral agent.

**Fig. 3.**
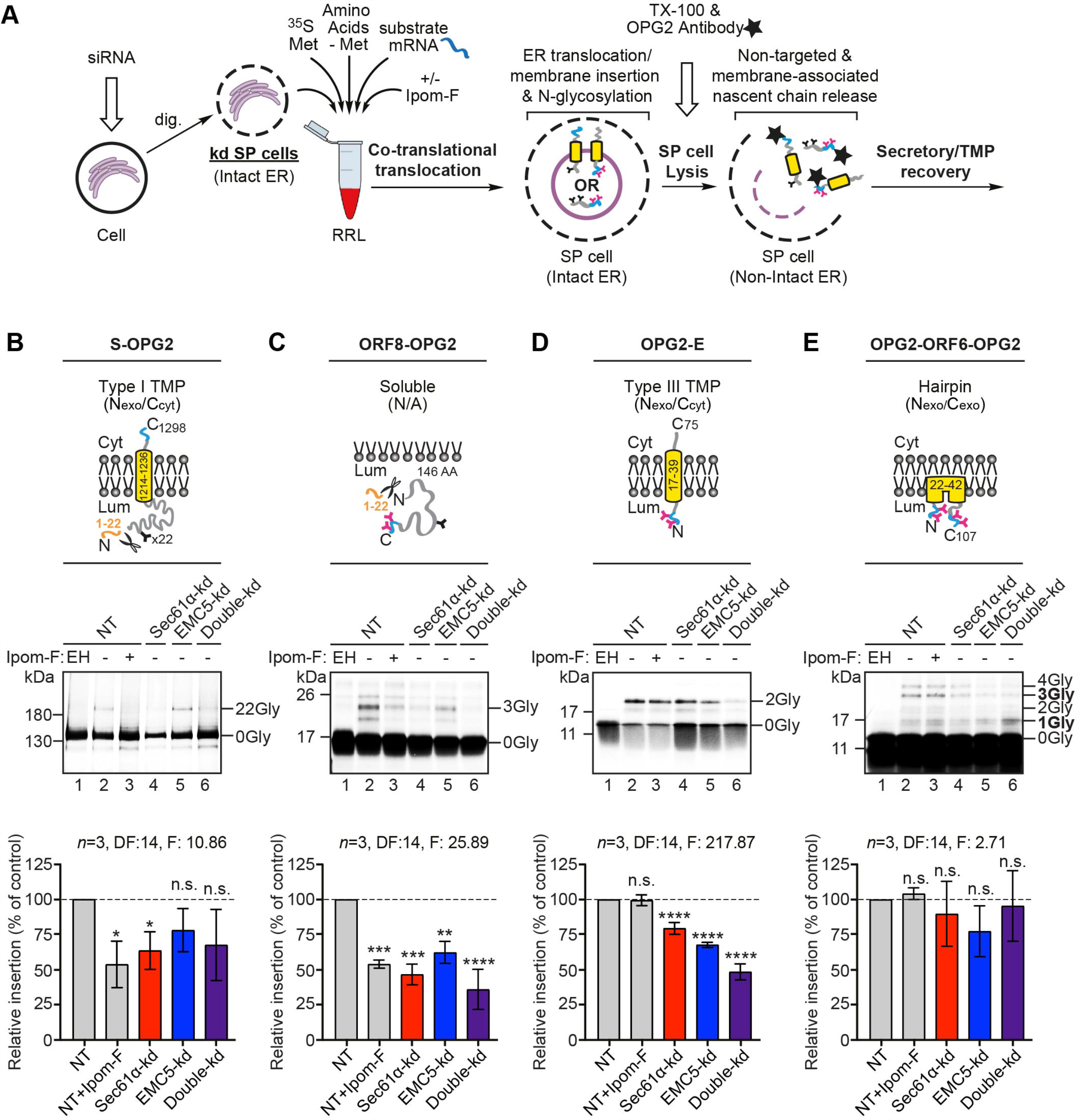
SARS-CoV-2 proteins are variably dependent on the Sec61 complex and/or the EMC for ER membrane translocation/insertion. **(A)** Schematic of *in vitro* ER import assay using control SP cells, or those depleted of a subunit of the Sec61 complex and/or the EMC via siRNA. Following translation, OPG2-tagged translation products (i.e. membrane-associated and non-targeted nascent chains) were immunoprecipitated, resolved by SDS-PAGE and analysed by phosphorimaging. OPG2-tagged variants of the SARS-CoV-2 **(B)** spike (S-OPG2), **(C)** ORF8 (ORF8-OPG2), **(D)** envelope (OPG2-E) and **(E)** ORF6 (OPG2-ORF6-OPG2 species (labelled as for Fig. 2) were synthesised in rabbit reticulocyte lysate supplemented with control SP cells (lanes 1-2) or those with impaired Sec61 and/or EMC function (lanes 3-6). Radiolabelled products were recovered and analysed as in **(A)**. Membrane translocation/insertion efficiency was determined using the ratio of the N-glycosylation of lumenal domains, identified using Endo H (EH, lane 1), relative to the NT control (set to 100% translocation/insertion efficiency). Quantitations (*n*=3) and statistical significance (two-way ANOVA, DF and F values shown in the figure) determined as for Figure 2. Statistical significance: n.s., non-significant >0.1; *, P < 0.05; **, P < 0.01; ***, P < 0.001; ****, P < 0.0001.

## Results and Discussion

### Ipom-F selectively inhibits ER translocation of the viral ORF8 and S proteins

To explore the ability of Ipom-F to inhibit the ER translocation of a small, yet structurally diverse, panel of SARS-CoV-2 membrane and secretory-like proteins, we first used a well-established *in vitro* translation system supplemented with canine pancreatic microsomes (Fig. 2A). To facilitate the detection of ER translocation, we modified the viral ORF8, S, E, M and ORF6 proteins by adding an OPG2-tag; an epitope that supports efficient ER lumenal N-glycosylation and enables product recovery via immunoprecipitation, without affecting Ipom-F sensitivity (Fig. S1A) (O’Keefe et al., 2020 submitted). For viral proteins that lack endogenous sites for N-glycosylation, such as the E protein, the ER lumenal OPG2-tag acts as a reporter for ER translocation and enables their recovery of by immunoprecipitation. Where viral proteins already contain suitable sites for N-glycosylation (S and M proteins), the cytosolic OPG2-tag is used solely for immunoprecipitation. The identity of the resulting N-glycosylated species for each of these OPG2-tagged viral proteins was confirmed by endoglycosidase H (Endo H) treatment of the radiolabelled products associated with the membrane fraction prior to SDS-PAGE (Fig. 2B, cf. lanes 1 and 2 in each panel).

Using ER lumenal modification of either endogenous N-glycosylation sites (viral S and M proteins) or the appended OPG2-tag (viral E and ORF8 proteins) as a reporter for ER membrane translocation, we found that 1 µM Ipom-F strongly inhibited both the translocation of the soluble, secretory-like protein ORF8-OPG2 and the integration of the type I transmembrane proteins (TMP) S-OPG2, and truncated derivatives thereof (Fig. 2B, Fig. 2C, Fig. S1C). Furthermore, membrane insertion of the human type I TMP, ACE2, was inhibited to a similar extent (Fig. 2B, Fig. 2C, ∼70 to ∼90% inhibition for these three proteins). These results mirror previous findings showing that precursor proteins bearing N-terminal signal peptides, and which are therefore obligate clients for the Sec61-translocon, are typically very sensitive to Ipom-F-mediated inhibition (Zong et al., 2019; O’Keefe et al., 2020 submitted). In the context of SARS-CoV-2 infection, wherein ACE2 acts as an important host cell receptor for the SARS-CoV-2 virus via its interaction with the viral S protein (Walls et al., 2020), these data suggest that an Ipom-F-induced antiviral effect might be achieved via selective reductions in the biogenesis of both host and viral proteins (cf. Fig. 1A).

In contrast to the viral S and ORF8 proteins, insertion of the viral E protein was unaffected by Ipom-F (Fig. 2B-C), consistent with its recent classification as a type III TMP (Duart et al., 2020). Type III TMP integration is highly resistant to Ipom-F (Zong et al., 2019), most likely because they exploit a novel pathway for ER insertion (cf. Fig. 3; O’Keefe et al., 2020 submitted). We therefore conclude that the known substrate-selective inhibitory action of Ipom-F at the Sec61 translocon is directly applicable to viral membrane proteins; whereby the ER translocation of secretory proteins and type I TMPs, but not type III TMPs, is efficiently blocked by Ipom-F.

The viral M protein is a multi-pass TMP with its first TMD oriented so the N-terminus is exoplasmic (N_exo_) and hence can be considered “type III-like”. Although human multi-pass TMPs of this type typically require both the ER membrane complex (EMC) and Sec61 translocon for their authentic ER insertion (Chitwood et al., 2018), Ipom-F had no significant effect on the ER translocation/insertion of the M protein *in vitro*, as judged by the efficiency of N-glycosylation of its N-terminal domain (Fig. 2C). We conclude that the integration of its first TMD is unaffected by Ipom-F, consistent with its use of the EMC (Chitwood et al., 2018; O’Keefe et al., 2020 submitted). There is however a qualitative reduction in the intensity of both the non- and N-glycosylated forms of the M protein when compared to the control (see Fig. 2B and Fig. S1A). We speculate that this decrease may reflect an Ipom-F-induced effect on the Sec61-dependent integration of the second and/or third TM-spans of the M protein (cf. Chitwood et al., 2018) and our future studies will aim to resolve this question. Nevertheless, like similar host cell multi-pass TMPs that are resistant to a similar Sec61 inhibitor mycolactone (Morel et al., 2018), the M protein appears more resistant to Ipom-F than either the S or ORF8 proteins (Fig 2, Fig. S1A). In practice, the potential resistance of this highly abundant and functionally diverse class of endogenous multi-spanning membrane proteins (von Heijne, 2007) may limit any Ipom-F-induced cytotoxicity towards host cells.

### ORF6 assumes a lumenal-facing hairpin topology in ER-derived microsomes

Cell-based studies of the ORF6 protein from SARS-CoV-1 suggest it has an unusual hairpin topology with both its N- and C-termini located on the exoplasmic side of the host cell membrane, to which it binds via an N-terminal amphipathic helix (Netland et al., 2007). To independently determine the membrane topology of SARS-CoV-2 ORF6, we prepared versions with OPG2 tags at both its N- and C-termini, or single tagged equivalents (see Fig. 2B, schematics, OPG2-ORF6-OPG2, OPG2-ORF6 and ORF6-OPG2). Following membrane insertion, doubly tagged OPG2-ORF6-OPG2 shows significant amounts of species with 3- and 4-N-linked glycans (Fig. 2B). This pattern confirms that the SARS-CoV-2 ORF6 protein assumes a ‘hairpin’ conformation in the ER membrane with both its N- and C-termini in the lumen (Fig. 2B, OPG2-ORF6-OPG2). These 3- and 4-N-glycan bearing OPG2-ORF6-OPG2 species are also resistant to extraction with alkaline sodium carbonate buffer (Fig. S1D) and protected from added protease (Fig. S1E), further indicating that the majority of the ORF6 protein is stably associated with the ER membrane in a ‘hairpin’ (N_exo_/C_exo_) topology.

Consistent with this unusual membrane topology, we find no indication that the membrane insertion of any of our OPG2-tagged ORF6 variants is reduced by Ipom-F, strongly suggesting that its association with the inner leaflet of the ER membrane does not require protein translocation via the central channel of the Sec61 translocon (Gérard et al., 2020; O’Keefe et al., 2020 submitted). We noted a sub-set of OPG2-ORF6-OPG2 species bearing only a single N-glycan was also clearly present in the membrane-associated fraction with or without Ipom-F treatment (Fig. 2B, OPG2-ORF6-OPG2, see 1Gly). Based on comparison to singly OPG2-tagged variants (Fig. 2B), we conclude that OPG2-ORF6-OPG2-1Gly has its N-terminus in the ER lumen, where only one of its two consensus sites is efficiently N-glycosylated (cf. Nilsson and von Heijne, 1993), whilst its C-terminus is either ER luminal but non-glycosylated or remains on the cytosolic side of the membrane. In the latter case, it may be that, in addition to its hairpin topology, some fraction of ORF6 may be integrated into ER-derived microsomes as a type III TMP (cf. Fig. S2E; see also Netland et al., 2007) that is resistant to Ipom-F inhibition (this study; Zong et al., 2019; O’Keefe et al., 2020 submitted).

### The molecular basis for SARS-CoV-2 protein sensitivity to Ipom-F

Having ascertained that Ipom-F inhibits the ER membrane translocation/insertion of the viral ORF8 and S proteins, but not of the ORF6, E or M proteins (Fig. 2C), we next investigated the molecular basis for this selectivity. For these studies we employed semi-permeabilised (SP) mammalian cells, depleted of specific membrane components via siRNA-mediated knockdown, as our source of ER membrane (Fig. 3A; Wilson et al., 2007). Consistent with our recent work (O’Keefe et al., 2020 submitted), and based on the quantitative immunoblotting of target and control gene products (Fig. S2A-C), we selectively depleted HeLa cells for core components of the Sec61 translocon (Sec61α-kd, ∼65% reduction), the EMC (EMC5-kd, ∼73% reduction) and both together (Sec61α+EMC5-kd, ∼68% and ∼78% reduction) prior to semi-permeabilisation with digitonin and use for *in vitro* ER translocation assays.

Following the analysis of total OPG2-tagged translation products recovered by immunoprecipitation, we found that: i) the S protein and a truncated derivative were both more strongly affected by the depletion of Sec61α than of EMC5 (Fig. 3B, Fig. S2D); ii) the ORF8 protein was likewise strongly affected by Sec61α depletion but also sensitive to EMC5 depletion (Fig. 3C); iii) the E protein showed diminished insertion efficiency after knock-down of Sec61α and EMC5, although the latter had a more pronounced effect (Fig. 3D). In each case, the combined knockdown of Sec61α and EMC5 resulted in a reduction of membrane insertion that was either comparable to, or greater than, that achieved following the knock-down of Sec61α alone (Figs. 3B to 3D). For the ORF6 protein, the total level of N-glycosylated OPG2-ORF6-OPG2 species was unaffected by any knockdown condition tested (Fig. 3E). However, we note a marked increase in the proportion of potentially mis-inserted OPG2-ORF6-OPG2-1Gly species, particularly after co-depletion of EMC5 and Sec61α (see Fig. 3E; Fig. S2E). We speculate that the unusual hairpin topology of the ORF6 protein may be attributed to the EMC and Sec61 complex acting in concert to provide an Ipom-F insensitive pathway for protein translocation across the ER membrane (O’Keefe et al., 2020, submitted). Perturbation of this pathway seemingly increases the potential for ORF6 to mis-insert (cf. Chitwood et al., 2018), perhaps as a consequence of disruption to the translocation of its C-terminus (Fig. S2E).

Taken together, our data establish that, analogous to human membrane and secretory proteins, the principal molecular basis for the Ipom-F-sensitivity of the SARS-CoV-2 ORF8 and S proteins is their dependence on Sec61-mediated protein translocation into and across the ER membrane. In contrast, the E, M, and ORF6 proteins appear capable of exploiting one or more alternative membrane insertion/translocation pathways that can bypass the translocase activity of the Sec61 complex. These alternatives most likely include a recently described route for type III TMP insertion that requires the insertase function of the EMC (O’Keefe et al., 2020 submitted), which our data suggest is also sufficient to confer Ipom-F-resistance to the viral E protein and at least the first TM-span of the viral M protein.

### Concluding Remarks

We conclude, that Sec61-selective protein translocation inhibitors like Ipom-F hold promise as broad-spectrum antivirals that may exert a therapeutic effect by selectively inhibiting the ER translocation of viral and/or host proteins which are crucial to viral infection and propagation (Mast et al., 2020). In the context of SARS-CoV-2, integration of the viral S protein and its host cell receptor, ACE2, into the ER membrane is significantly reduced by Ipom-F (Fig. 2C, 3B). Likewise, translocation of the viral ORF8 protein across the ER membrane and into its lumen is substantially diminished (Fig. 2C, 3C). The binding of the viral S protein to cell surface ACE2 is a key step in host cell infection (Drew and Janes, 2020), whilst ORF8 may protect SARS-CoV-2 infected cells against host cytotoxic T lymphocytes (Zhang et al., 2020), making all three of these proteins viable therapeutic targets (Drew and Janes, 2020; Li et al., 2020; Young et al., 2020).

Like other small molecule inhibitors that target fundamental cellular pathways (Bojkova et al. 2020), the broad-ranging effects of Sec61 inhibitors on host cell membrane and secretory protein synthesis (Morel et al., 2018; Zong et al. 2019), including the strong *in vitro* effect of Ipom-F on ACE2 biogenesis (cf. Groβ et al. 2020), present an obvious hurdle to their future use. Nevertheless, given that Ipom-F is a potent inhibitor of Sec61-mediated protein translocation in cell culture models (Zong et al., 2019), and appears well tolerated in mice (Zong et al., 2020), we propose that future studies investigating its effect on SARS-CoV-2 infection and propagation in cellular models are clearly warranted (cf. Bojkova et al. 2020).

## Materials and Methods

### Ipom-F and Antibodies

Ipom-F was synthesised as previously described (Zong et al., in press). Antibodies used to validate Sec61 and/or EMC subunit depletions in SP cells (Fig. S2) were purchased from Santa Cruz Biotechnology (goat polyclonal anti-LMNB1 (clone M-20, sc-6217), Bethyl Laboratories (rabbit polyclonal anti-EMC5 (A305-832-A)), Abcam (rabbit polyclonal anti-EMC6, (ab84902)), gifted by Sven Lang and Richard Zimmermann (University of Saarland, Homburg, Germany, rabbit anti-Sec61α) or as previously described (mouse monoclonal anti-OPG2 tag (McKenna et al., 2016) and rabbit polyclonal anti-OST48 (Wilson et al., 2007).

### DNA constructs

The cDNA for human ACE2 (Uniprot: Q9BYF1) was purchased from Sino Biological (HG10108-M). cDNAs encoding the SARS-CoV-2 genes for ORF6, ORF8 and the E M and S proteins (Uniprot: P0DTC6, P0DTC8, P0DTC4, P0DTC5, P0DTC2 respectively) were kindly provided by Nevan Krogan (UCSF, US) (Gordon et al. 2020), amplified by PCR, subcloned into the pcDNA5 vector and constructs validated by DNA sequencing (GATC, Eurofins Genomics). ORF6-OPG2, ORF8-OPG2, M-OPG2 and S-OPG2 were generated by inserting the respective cDNAs in frame between NheI and AflII sites of a pcDNA5/FRT/V5-His vector (Invitrogen) containing a C-terminal OPG2 tag (MNGTEGPNFYVPFSNKTG). OPG2-E was generated by cloning the cDNA encoding the E-protein into the same pcDNA5-OPG2 vector using the KpnI and BamHI sites and deleting the stop codon after the OPG2 tag by site-directed mutagenesis (Stratagene QuikChange, Agilent Technologies). The N-terminal OPG2-tag of OPG2-ORF6-OPG2 was inserted by site-directed mutagenesis of ORF6-OPG2 using the relevant forward and reverse primers (Integrated DNA Technologies). Linear DNA templates were generated by PCR and mRNA transcribed using T7 polymerase.

### siRNA-mediated knockdown and SP cell preparation

HeLa cells (human epithelial cervix carcinoma cells) were cultured in DMEM supplemented with 10% (v/v) FBS and maintained in a 5% CO_2_ humidified incubator at 37°C. Knockdown of target genes were performed as previously described (O’Keefe et al., 2020 submitted) using 20 nM (final concentration) of either control siRNA (ON-TARGETplus Non-targeting control pool; Dharmacon), *SEC61A1* siRNA (Sec61α-kd, GE Healthcare, sequence AACACUGAAAUGUCUACGUUUUU), *MMGT1* siRNA (EMC5-kd, ThermoFisher Scientific, s41129) and INTERFERin (Polyplus, 409-10) as described by the manufacturer. 96 h post-initial transfection, cells were semi-permeabilsed using 80 μg/mL high purity digitonin (Calbiochem) and treated with 0.2 U Nuclease S7 Micrococcal nuclease from *Staphylococcus aureus* (Sigma-Aldrich, 10107921001) as previously described (O’Keefe et al., 2020 submitted; Wilson et al., 2007). SP cells lacking endogenous mRNA were resuspended (3×10^6^ SP cells/mL as determined by trypan blue (Sigma-Aldrich, T8154) staining) in KHM buffer (110 mM KOAc, 2 mM Mg(OAc)_2_, 20 mM HEPES-KOH pH 7.2) prior to analysis by western blot, or inclusion in translation master mixes such that each translation reaction contained 2×10^5^ cells/mL.

### *In vitro* ER import assays

Standard translation and membrane translocation/insertion assays, supplemented with nuclease-treated canine pancreatic microsomes (from stock with OD_280_ = 44/mL) or siRNA-treated SP HeLa cells, were performed in nuclease-treated rabbit reticulocyte lysate (Promega) as previously described (Zong et al., 2019; O’Keefe et al., 2020 submitted): namely in the presence of EasyTag EXPRESS ^35^S Protein Labelling Mix containing [^35^S] methionine (Perkin Elmer) (0.533 MBq; 30.15 TBq/mmol), 25 μM amino acids minus methionine (Promega), 1 μM Ipom-F, or an equivalent volume of DMSO, 6.5% (v/v) ER-derived microsomes or SP cells and ∼10% (v/v) of *in vitro* transcribed mRNA (∼500 ng/μL) encoding the relevant precursor protein. Microsomal translation reactions (20 μL) were performed for 30 min at 30°C whereas those using SP HeLa cells were performed on a 1.5X scale (30 μL translation reactions) for 1 h at 30°C. As the S protein was most efficiently synthesised using the TNT® Coupled system (Fig. S1B), import assays of the comparatively higher molecular weight ACE2 and S proteins (50 μL reactions) were both performed using the TNT® Coupled Transcription/ Translation system (Promega) for 90 min at 30°C as described by the manufacturer (∼50 ng/μL cDNA,1 μM Ipom-F or an equivalent volume of DMSO, 12% (v/v) ER-derived microsomes or SP cells). All translation reactions were finished by incubating with 0.1 mM puromycin for 10 min at 30°C to ensure translation termination and ribosome release of newly synthesised proteins prior to analysis.

### Recovery and analysis of radiolabelled products

Following puromycin treatment, microsomal membrane-associated fractions were recovered by centrifugation through an 80 μL high-salt cushion (0.75 M sucrose, 0.5 M KOAc, 5 mM Mg(OAc)_2_, 50 mM Hepes-KOH, pH 7.9) at 100,000 ***g*** for 10 min at 4°C and the pellet suspended directly in SDS sample buffer. To confirm the topology of ORF6 (Fig. S2), the membrane-associated fraction of the doubly-OPG2-tagged form (OPG2-ORF6-OPG2) was resuspended in KHM buffer (20 μL) and subjected to either carbonate extraction (0.1 M Na_2_CO_3_, pH 11.3) (McKenna et al., 2016) or a protease protection assay using trypsin (1 μg/mL) with or without 0.1% Triton X-100 (Ray-Sinha et al., 2009) prior to suspension in SDS sample buffer. For translation reactions using SP cells, the total reaction material was diluted with nine volumes of Triton immunoprecipitation buffer (10 mM Tris-HCl, 140 mM NaCl, 1 mM EDTA, 1% Triton X-100, 5 mM PMSF, 1 mM methionine (to prevent background from the radiolabelled methionine), pH 7.5). Samples were incubated under constant agitation with an antibody recognising the OPG2 epitope (1:200 dilution) for 16 h at 4°C to recover both the membrane-associated and non-targeted nascent chains. Samples were next incubated under constant agitation with 10% (v/v) Protein-A-Sepharose beads (Genscript) for a further 2 h at 4°C before recovery by centrifugation at 13,000 ***g*** for 1 min. Protein-A-Sepharose beads were washed twice with Triton immunoprecipitation buffer prior to suspension in SDS sample buffer. Where indicated, samples were treated with 1000 U of a form of Endoglycosidase H that does not co-migrate with and hence potentially distort the radiolabelled products when resolved: Endoglycosidase Hf (translation products of ∼10-50 kDa; New England Biolabs, P0703S) or Endoglycosidase H (translation products of ∼50-150 kDa protein substrates; New England Biolabs, P0702S). All samples were solubilised for 12 h at 37°C and then sonicated prior to resolution by SDS-PAGE (10% or 16% PAGE, 120V, 120-180 min). Gels were fixed for 5 min (20% MeOH, 10% AcOH), dried for 2 h at 65°C and radiolabelled products visualised using a Typhoon FLA-700 (GE Healthcare) following exposure to a phosphorimaging plate for 24-72 h.

### Western Blotting

Following semi-permeabilisation, aliquots of siRNA-treated HeLa cells were suspended in SDS sample buffer, denatured for 12 h at 37°C and sonicated prior to resolution by SDS-PAGE (16% or 10% PAGE, 120V, 120-150 min). Following transfer to a PVDF membrane in transfer buffer (0.06 M Tris, 0.60 M glycine, 20% MeOH) at 300 mA for 2.5 h, PVDF membranes were incubated in 1X Casein blocking buffer (10X stock from Sigma-Aldrich, B6429) made up in TBS, incubated with appropriate primary antibodies (1:500 or 1:1000 dilution) and processed for fluorescence-based detection as described by LI-COR Biosciences using appropriate secondary antibodies (IRDye 680RD Donkey anti-Goat, IRDye 680RD Donkey anti-Rabbit, IRDye 800CW Donkey anti-Mouse) at 1:10,000 dilution. Signals were visualised using an Odyssey CLx Imaging System (LI-COR Biosciences).

### Quantitation and Statistical Analysis

Bar graphs depict either the efficiency of membrane translocation/insertion calculated as the ratio of N-glycosylated protein relative to the amount of non-N-glycosylated protein (Fig. 2-3), or the efficiencies of siRNA-mediated knockdown in SP cells calculated as a proportion of the protein content when compared to the NT control (Fig. S2), with all control samples set to 100%. Normalised values were used for statistical comparison (one-way or two-way ANOVA; DF and F values are shown in each figure as appropriate and the multiple comparisons test used are indicated in the appropriate figure legend). Statistical significance is given as n.s., non-significant >0.1; *, P < 0.05; **, P < 0.01; ***, P < 0.001; ****, P < 0.0001.

## Acknowledgements

We thank Quentin Roebuck for technical assistance, Nevan Krogan (UCSF) for SARS-CoV-2 plasmids, Sven Lang (University of Saarland) for Sec61α antisera, Belinda Hall and Rachel Simmonds (University of Surrey) for useful discussions. We are indebted to Richard Zimmermann (University of Saarland) for catalyzing SARS-CoV-2 related discussions amongst the ER research community.

## Author Contributions

K.B.D., G.Z. and H.S. participated in synthesis of Ipom-F and W.Q.S supervised the synthesis; P.R. generated SARS-CoV-2 plasmids; S.O’K. performed site-directed mutagenesis and experiments; S.O’K. and S.H. designed the study, analysed the data and wrote the manuscript.

## Supplementary Information

Supplementary information Fig. S1 and Fig. S2 accompanies this report.

## Supplementary information

**Fig. S1.**
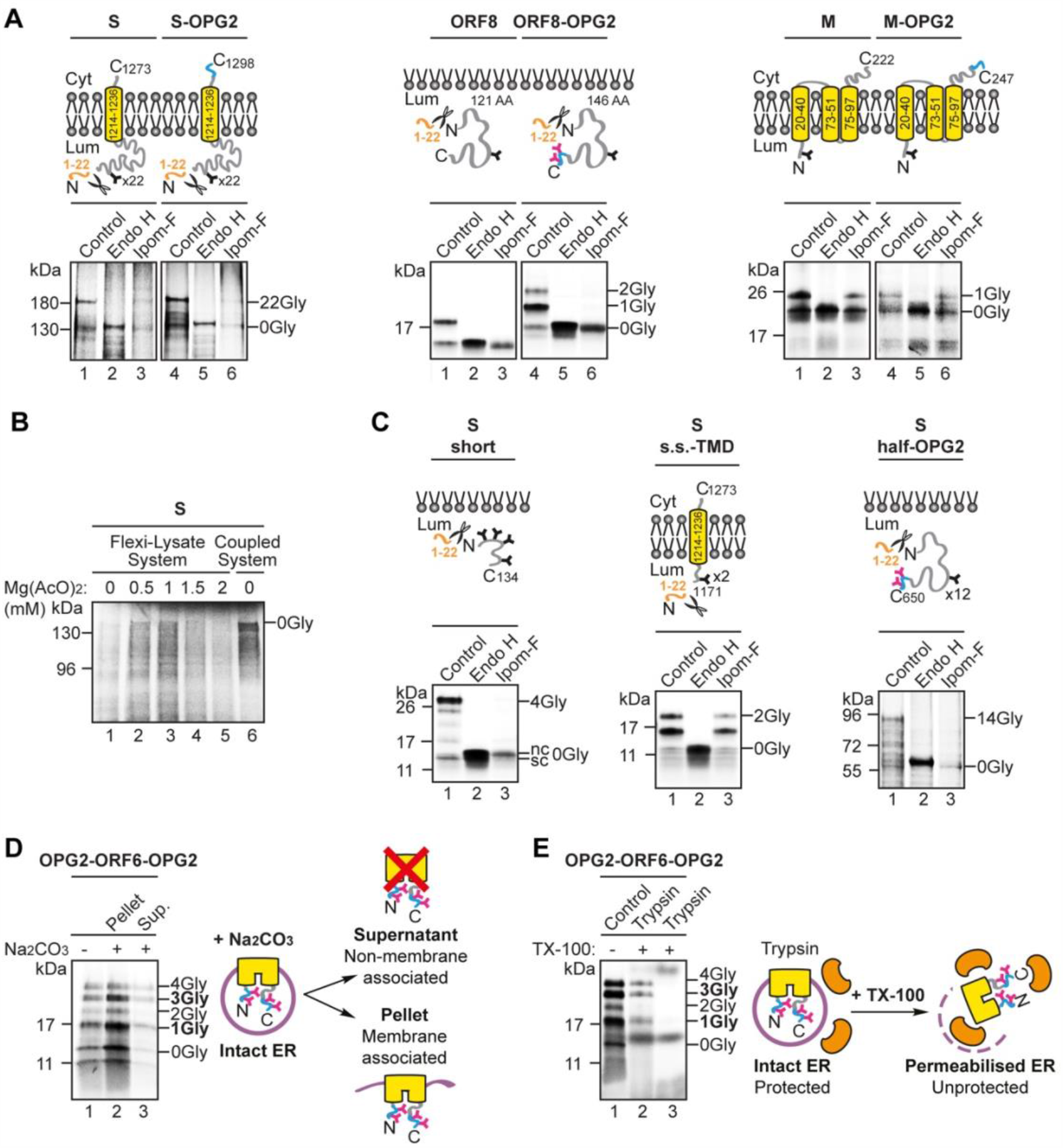
Additional studies using ER microsomes, Related to Figures 2 and 3. (**A**) Non-tagged (lanes 1-3) and OPG2-tagged (lanes 4-6) versions of the SARS-CoV-2 spike protein (S, S-OPG2), ORF8 (ORF8, ORF8-OPG2) and membrane protein (M, M-OPG2) were synthesised in rabbit reticulocyte lysate supplemented with ER-derived canine pancreatic microsomes in the absence and presence of Ipom-F (lanes 1 and 3). Phosphorimages of membrane-associated products resolved by SDS-PAGE together with representative substrate outlines are shown. N-glycosylated (X-Gly) versus non-N-glycosylated (0Gly) species were identified by treatment with endoglycosidase H (Endo H, lanes 2 and 5). (**B**) The S protein was synthesised in a Flexi® rabbit reticulocyte system with varying concentrations of magnesium acetate (lanes 1-5) and a TNT® Coupled system (lane 6) in the absence of ER-derived microsomes. 5% of the total reaction material was resolved by SDS-PAGE and visualised by phosphorimaging. (**C**) The ER import of truncated variants of the S protein (S-short, S-s.s.-TMD, S-half-OPG2) was analysed as described for **(A)**. (**D**) The membrane-associated products of the doubly tagged form of ORF6 (OPG2-ORF6-OPG2) were synthesised as in **(A)** and, following treatment with sodium carbonate buffer and centrifugation, the pellet, enriched for membrane-integrated material, and supernatant, largely containing peripherally membrane-associated material, were analysed for OPG2-ORF6-OPG2. (**E**) The membrane-associated products of OPG2-ORF6-OPG2 were treated with trypsin in the absence or presence of Triton-100 (TX-100, lanes 2-3).

**Fig. S2.**
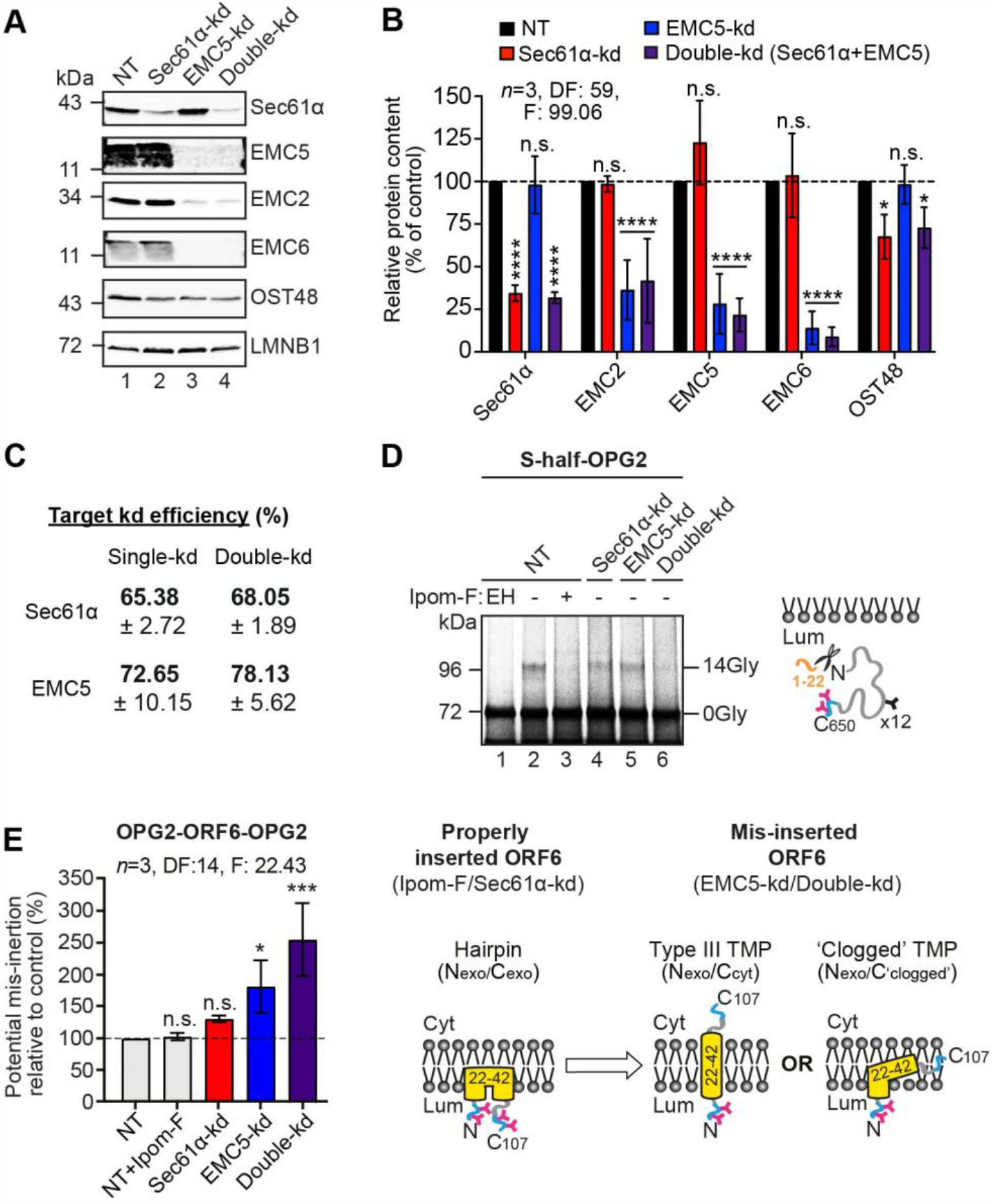
Validation of Sec61 and/or EMC subunit depletions in SP cells, Related to Figure 3. **(A)** The effects of transfecting HeLa cells with non-targeting (NT; lane 1), Sec61α-targeting (lane 2), EMC5-targeting (lane 3) and Sec61α+EMC5-targeting (lane 4) siRNAs were determined after semi-permeabilisation by immunoblotting for target genes (Sec61α, EMC5). Controls to assess destabilisation of the wider EMC complex (EMC2 and EMC6), any effect on the N-glycosylation machinery (the ER-resident 48 kDa subunit of the oligosaccharyl-transferase complex (OST48) and the quantity of SP cells used in each experiment (the nuclear protein Lamin-B1 (LMNB1)), are also shown. (**B**) The efficiencies of siRNA-mediated knockdown (bold) were calculated as a proportion of the signal intensity obtained with the NT control (set as 100%). Quantitations are given as mean±s.e.m for three separate siRNA treatments (*n*=3) with statistical significance of siRNA-mediated knockdowns (two-way ANOVA, DF and F values shown in the figure) determined using Dunnett’s multiple comparisons test. Statistical significance is given as n.s., non-significant >0.1; *, P < 0.05; ****, P < 0.0001. (**C**) Knockdown efficiencies (mean±s.e.m) for each of the target genes. (**D**) A truncated variant of the S protein (S-half-OPG2) was synthesised in rabbit reticulocyte lysate supplemented with SP cells with impaired Sec61 complex and/or EMC function and recovered by immunoprecipitation via the OPG2 tag. Radiolabelled products resolved by SDS-PAGE and analysed by phosphorimaging. N-glycosylated (14-Gly) versus non-N-glycosylated (0Gly) species were identified by treatment with endoglycosidase H (Endo H, lane 1). (**E**) Further analysis of the data presented in Fig. 3E of the main text. Here, the ratio of 3Gly and 4Gly bearing OPG2-ORF6-OPG2 N-glycosylated species relative to the 1Gly species present in the same sample was used as a proxy to estimate potential mis-insertion of the ORF6 protein in SP cells with impaired Sec61 complex and/or EMC function relative to the NT control (set to 100% efficiency). Quantitations are given as mean±s.e.m for independent translation reactions from separate siRNA treatments performed in triplicate (*n*=3) and statistical significance (two-way ANOVA, DF and F values shown in the figure) was determined using Dunnett’s multiple comparisons test. Statistical significance is given as n.s., non-significant >0.1; *, P < 0.05; ***, P < 0.001.

